# Essential roles of HGT-derived sterol biosynthesis in trypanosomatid growth and parasitism

**DOI:** 10.1101/2025.08.28.672796

**Authors:** Xuye Yuan, Tatsuhiko Kadowaki

## Abstract

Horizontal gene transfer (HGT) has played a major role in the evolution of kinetoplastids, but its scope and functional significance remain incompletely understood. We performed a comparative genomic analysis of 12 trypanosomatid parasites together with the free-living kinetoplastid *Bodo saltans* as an outgroup, using the DarkHorse algorithm to identify putative HGT candidates. *B. saltans* encoded the largest number of HGT-derived genes (980; 5.2% of coding sequences), whereas parasitic *Leishmania* species retained the fewest (67-71; 0.8-0.9%), consistent with extensive gene loss accompanying the transition to parasitism. Dollo parsimony reconstruction indicated that the kinetoplastid ancestor acquired numerous horizontally transferred genes, many of which were subsequently lost, but several functionally important families were retained in trypanosomatids. Among these, we identified a bacterial-derived aspartate ammonia ligase that, together with aspartate ammonia lyase, likely supports redox balance and succinate fermentation under anaerobic conditions in the insect gut. Phylogenetic analyses also revealed bacterial or archaeal origins of multiple enzymes in ergosterol biosynthesis, an essential pathway in trypanosomatids. Functional characterization in *Lotmaria passim* demonstrated that sterol C-8 isomerase is indispensable, whereas sterol C-5 desaturase is nonessential *in vitro* but critical for parasite growth at low temperature, amphotericin B sensitivity, and successful colonization of the honey bee hindgut. These findings illustrate the dual evolutionary trajectory of HGT in kinetoplastids: extensive gene loss during the transition to parasitism, coupled with the retention of lineage-specific innovations essential for survival in host environments.

**Significance Statement:** Kinetoplastids include both free-living species such as *Bodo saltans* and parasitic trypanosomatids, the latter responsible for devastating human, animal, and insect diseases. While trypanosomatids have streamlined their genomes during host adaptation, free-living relatives retained a broader genetic repertoire shaped by environmental gene acquisition. By systematically tracing horizontally transferred genes (HGTs) across kinetoplastids, we reveal how free-living *B. saltans* accumulated diverse HGT candidates, whereas parasitic trypanosomatids lost many of these genes but selectively retained or acquired others critical for their specialized niches. We show that HGT contributed to the evolution of sterol biosynthesis and metabolic pathways supporting parasite survival in anaerobic insect guts. Functional analyses in *Lotmaria passim*, a honey bee parasite, demonstrate that HGT-dependent ergosterol biosynthesis is essential for gut colonization and influence sensitivity to amphotericin B. These findings highlight how horizontal gene transfer not only shaped the metabolic architecture of kinetoplastids but also underpins parasite adaptation to host environments.

## Introduction

Kinetoplastids are a diverse group of flagellated protists defined by the presence of a kinetoplast, a unique mitochondrial DNA network. They include both free-living taxa, such as *Bodo saltans*, and parasitic lineages within the family Trypanosomatidae, which infect a wide range of invertebrate and vertebrate hosts [1–3]. Parasitic trypanosomatids include medically and agriculturally important pathogens such as *Trypanosoma brucei*, the causative agent of African sleeping sickness; *Trypanosoma cruzi*, the agent of Chagas disease; and *Leishmania* spp., which cause leishmaniasis in humans [4–6]. Other trypanosomatids, including *Lotmaria passim* and *Crithidia bombi*, are gut parasites of insects, with emerging ecological importance in pollinator health [7, 8].

The evolutionary transition from a free-living to parasitic lifestyle was accompanied by major shifts in genome content and metabolic capacity. Comparative genomics has shown that trypanosomatids exhibit streamlined genomes relative to bodonids such as *B. saltans*, consistent with large-scale gene loss during adaptation to host-associated niches [3, 9]. Nevertheless, trypanosomatids have also acquired lineage-specific innovations, some of which appear to derive from horizontal gene transfer (HGT). HGT is recognized as an important evolutionary force in eukaryotes, especially in microbial lineages, where it contributes to metabolic novelty, ecological adaptation, and host–parasite interactions [10–12]. Evidence for HGT in kinetoplastids has accumulated from both phylogenomic analyses and targeted studies of individual metabolic enzymes. Early work identified bacterial-like enzymes localized to the glycosome in *Leishmania* and *Trypanosoma* species, suggesting acquisition from prokaryotes [13, 14]. More recently, systematic surveys have proposed that kinetoplastids inherited multiple bacterial genes involved in carbohydrate and amino acid metabolism [15]. However, the full extent, evolutionary trajectory, and functional relevance of HGT in kinetoplastids remain to be resolved.

One metabolic context in which HGT may be particularly significant is adaptation to the insect gut. Many trypanosomatids inhabit the hindgut of insects, an environment that is frequently anaerobic. In these conditions, trypanosomatids rely on succinate fermentation to sustain ATP production through glycolysis [16, 17]. Enzymes such as fumarate reductase are central to this process, and recent findings suggest that bacterial-derived genes may contribute additional pathways to balance redox equivalents under oxygen limitation [18].

Another context where HGT has been implicated is sterol biosynthesis. Unlike animals, which synthesize cholesterol, or plants, which produce phytosterols, kinetoplastids synthesize ergosterol, a sterol otherwise characteristic of fungi [19]. Ergosterol is essential for kinetoplastid membrane structure and is the molecular target of amphotericin B, a frontline antileishmanial drug [20]. Functional studies in *Leishmania* spp. have confirmed that disruption of sterol biosynthetic genes alters parasite viability, morphology, and drug susceptibility [21–23]. Yet, systematic integration of comparative genomics, phylogenetic analysis, and functional validation is still lacking.

In this study, we combined large-scale comparative genomics of 12 trypanosomatid species with phylogenetic and experimental analyses to assess the scope and functional significance of HGT in kinetoplastids. By including the free-living *B. saltans* as an outgroup, we reconstructed the evolutionary history of HGT-derived genes and distinguished ancestral acquisitions from lineage-specific innovations. We highlight two major findings: (i) the acquisition of bacterial-derived enzymes that may facilitate anaerobic metabolism in insect gut environments, and (ii) the ergosterol biosynthetic pathway with the HGT derived key enzymes to be essential for parasite growth at low temperature and honey bee gut colonization. Together, our results illustrate how HGT has shaped the evolution of kinetoplastids, coupling large-scale gene loss with the selective retention of metabolic innovations critical for parasitism.

## Results

### Identification of horizontally transferred genes in trypanosomatids

To identify candidate genes acquired by HGT, we applied the DarkHorse algorithm [24] to 12 trypanosomatid genomes, using the free-living kinetoplastid *B. saltans* as an outgroup. DarkHorse infers HGT candidates by evaluating the lineage probability index (LPI), which measures the phylogenetic relatedness of a gene relative to its best BLAST hits. Genes with LPI < 0.6 were classified as putative HGT candidates (Table S1). *B. saltans* harbored the highest number (980 genes; 5.2% of its coding repertoire), whereas parasitic *Leishmania* spp. encoded the fewest (67–71 genes; 0.8–0.9%). Other trypanosomatids, including *T. brucei, T. cruzi, Leptomonas seymouri, Leptomonas pyrrhocoris, C. bombi*, and *L. passim*, carried intermediate proportions (0.75–1.9%). This pattern supports the view that the transition from a free-living to parasitic lifestyle was accompanied by large-scale gene loss [3], including many HGT-derived genes.

### Evolutionary gain and loss of HGT-derived gene families

To trace the evolutionary history of HGT candidates, we grouped them into orthogroups using OrthoFinder and reconstructed ancestral states with Dollo parsimony (Fig. 1). *B. saltans* was predicted to have acquired 488 gene families; however, it is likely that most of them were subsequently lost in the common ancestor of parasitic trypanosomatids. Nonetheless, this ancestor gained 44 additional HGT-derived families after diverging from *B. saltans*. While isolated branches (e.g., the ancestors of *Trypanosoma* spp. and of Leishmaniinae) showed net gene family gains, losses were generally more frequent. These data suggest that acquisition of new genes by HGT is rare, and their retention depends strongly on lineage-specific ecological pressures.

**Figure 1.**
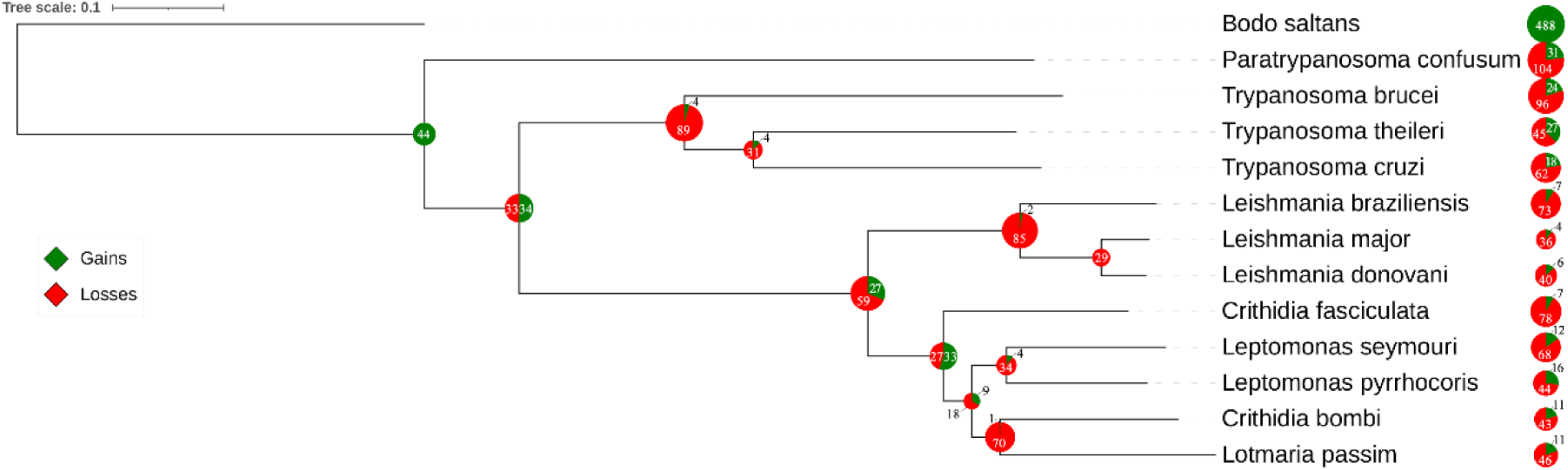
Gains and losses of HGT-derived gene families in kinetoplastids inferred by Dollo parsimony. The maximum likelihood tree of 13 kinetoplastid species was reconstructed based on previous reports. The horizontal bar indicates 0.1 substitutions per site. HGT candidates were grouped into orthogroups (OGs) using OrthoFinder, and the numbers of gene family gains (green) and losses (red) are shown for each branch and species.

### HGT and adaptation to anaerobic environments

Many trypanosomatids colonize insect hindguts, which are low in oxygen. We identified an aspartate ammonia ligase gene in trypanosomatids that is otherwise restricted to bacteria, as well as aspartate ammonia lyase clustering with bacterial homologs in phylogenetic analysis (Fig. 2). These enzymes establish a nitrogen–redox balancing cycle: aspartate ammonia lyase generates fumarate and ammonia, and aspartate ammonia ligase recaptures ammonia into asparagine, preventing accumulation. Fumarate can subsequently be reduced to succinate by fumarate reductase, regenerating NAD⁺ under anaerobic conditions [25]. This mechanism likely sustains glycosomal glycolysis and ATP production via substrate-level phosphorylation. Notably, the ligase is absent from the free-living *B. saltans*, suggesting that acquisition via HGT enabled adaptation to insect gut niches. Additional glycolytic enzymes previously proposed to be acquired by HGT [13, 16] reinforce this view.

**Figure 2.**
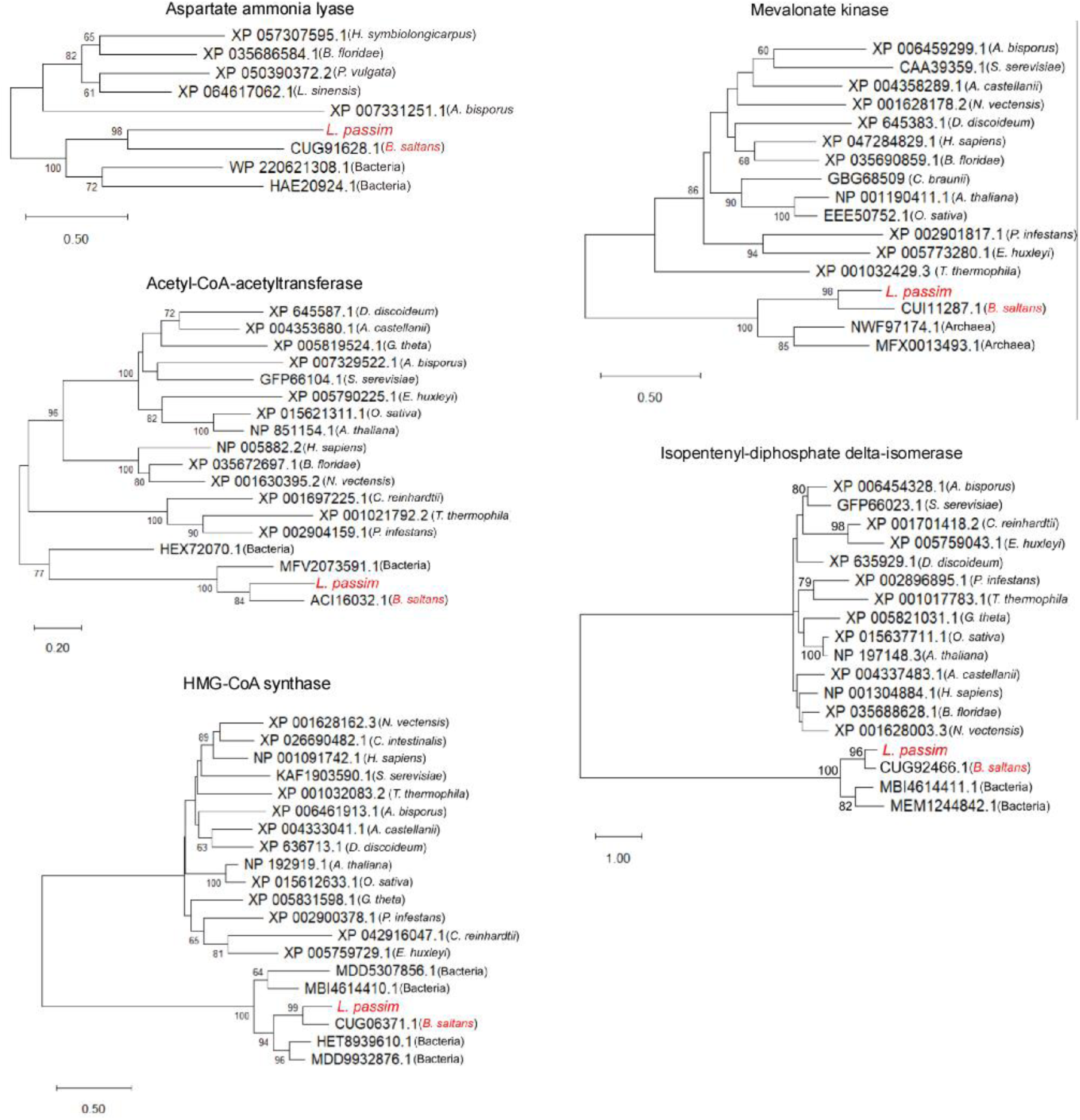
Evolutionary analysis of five HGT candidates using the maximum likelihood method. Phylogenetic trees were reconstructed for Aspartate ammonia lyase, Acetyl-CoA acetyltransferase, HMG-CoA synthase, Mevalonate kinase, and Isopentenyl-diphosphate delta-isomerase from *Lotmaria passim*, *Bodo saltans*, various eukaryotes, bacteria, and archaea using the maximum likelihood method. Protein sequences are labeled with accession numbers and species names. Bootstrap values above 60 are shown at the nodes. Horizontal bars indicate 0.2–1 substitutions per site, depending on the protein.

### HGT origins of ergosterol biosynthesis enzymes

Unlike animals and plants, which produce cholesterol or phytosterols, kinetoplastids and fungi synthesize ergosterol (Fig. 3). Early steps in sterol biosynthesis are conserved with bacteria, raising the possibility of bacterial origins. Phylogenetic analyses revealed that acetyl-CoA acetyltransferase, HMG-CoA synthase, and isopentenyl-diphosphate delta-isomerase from *L. passim* and *B. saltans* cluster with bacterial proteins, while mevalonate kinase groups with archaeal homologs (Fig. 2). These results indicate that multiple early enzymes in the sterol biosynthesis pathway were acquired by HGT in the kinetoplastid ancestor, consistent with prediction of glycosomal localization of mevalonate kinase and isopentenyl-diphosphate delta-isomerase in *Leishmania major* [26].

**Figure 3.**
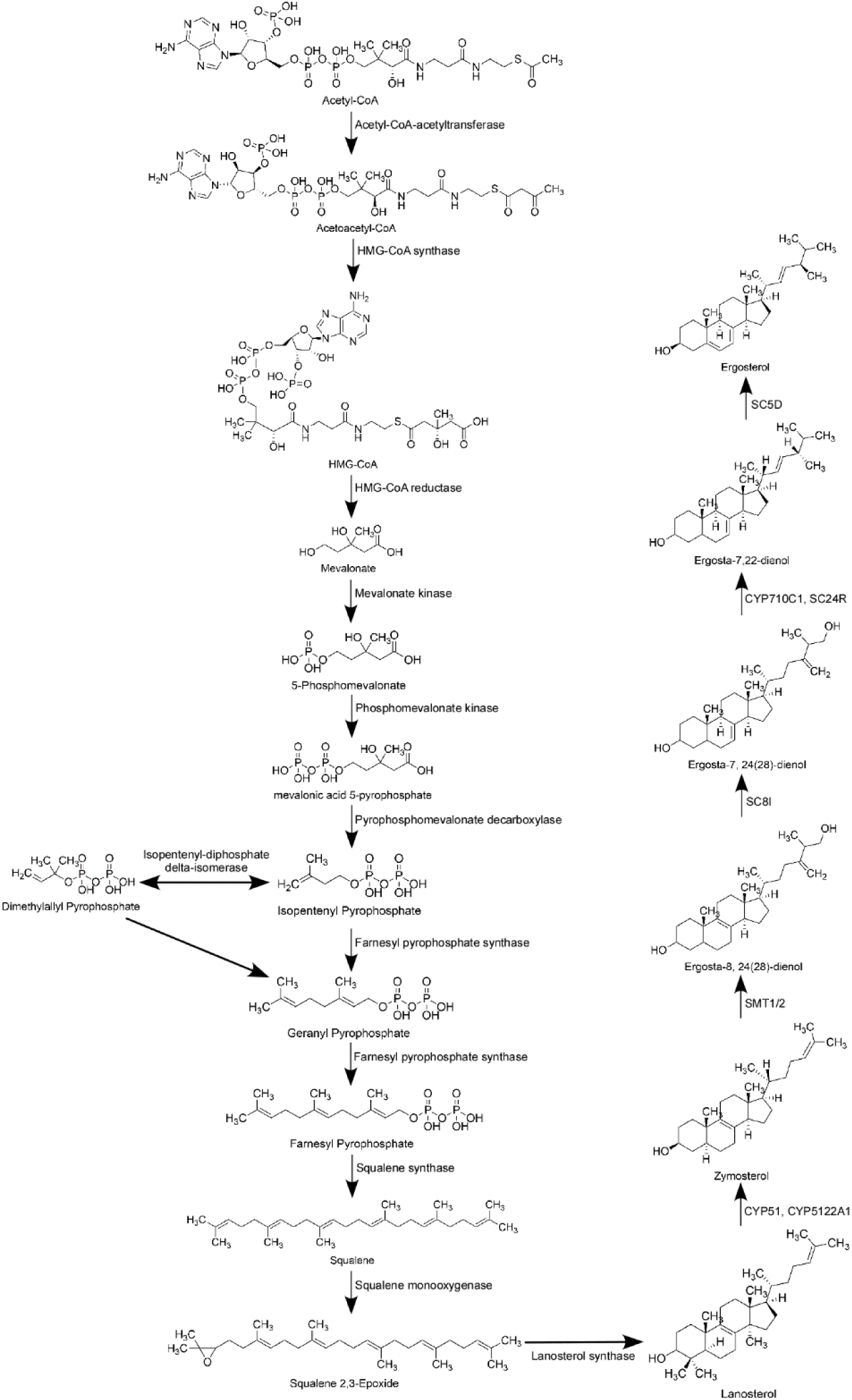
Ergosterol biosynthesis pathway. The pathway illustrates the metabolic steps from acetyl-CoA to ergosterol in trypanosomatids, along with the corresponding enzymes catalyzing each step. The pathway may branch from zymosterol if the enzymes act in a different order.

### Essentiality of sterol C-8 isomerase but not C-5 desaturase in *L. passim*

Because the four enzymes acquired by HGT participate in the early steps of ergosterol biosynthesis, their disruption is predicted to be lethal for *L. passim*. We thus targeted genes encoding sterol C-8 isomerase (LpSC8I) and sterol C-5 desaturase (LpSC5D), which act at the later stages of ergosterol synthesis (Fig. 3). The *L. passim* genome contains a single copy of *LpSC8I*, but two copies of *LpSC5D* (*LpSC5D1* and *LpSC5D2*). This duplication is also present in *L. pyrrhocoris, C. bombi*, and *Crithidia fasciculata*, but absent in *L. seymouri*, suggesting that *SC5D* duplicated in a common ancestor of the former five species and was subsequently lost in *L. seymouri.* We found that LpSC8I, LpSC5D1, and LpSC5D2 co-localize with the ER marker protein LpBiP (Fig. 4), suggesting that these enzymes are concentrated in the ER. Thus, ergosterol biosynthesis in trypanosomatids likely occurs in both the glycosome and the ER. CRISPR-based homology-directed repair was used to disrupt *LpSC8I* [27]. However, despite repeated attempts—including electroporation of a heterozygous knock-out clone with the knock-out construct—we failed to obtain homozygous mutants. This indicates that LpSC8I is essential for viability in *L. passim*, and that Ergosta-8,24(28)-dienol cannot substitute for ergosterol. In contrast, we successfully disrupted both *LpSC5D1* and *LpSC5D2* by replacing their open reading frames with the hygromycin phosphotransferase (*Hph*) gene (Fig. 5A). Diagnostic PCR confirmed the absence of wild-type alleles at the 5’ end of *LpSC5D1* and 3’ end of *LpSC5D2* (Fig. 5B). RT-PCR using a reverse primer complementary to both *LpSC5D1* and *LpSC5D2* confirmed that both transcripts were absent in homozygous mutants (Fig. 5C). Unlike LpSC8I, LpSC5D is not essential for viability under culture conditions.

**Figure 4.**
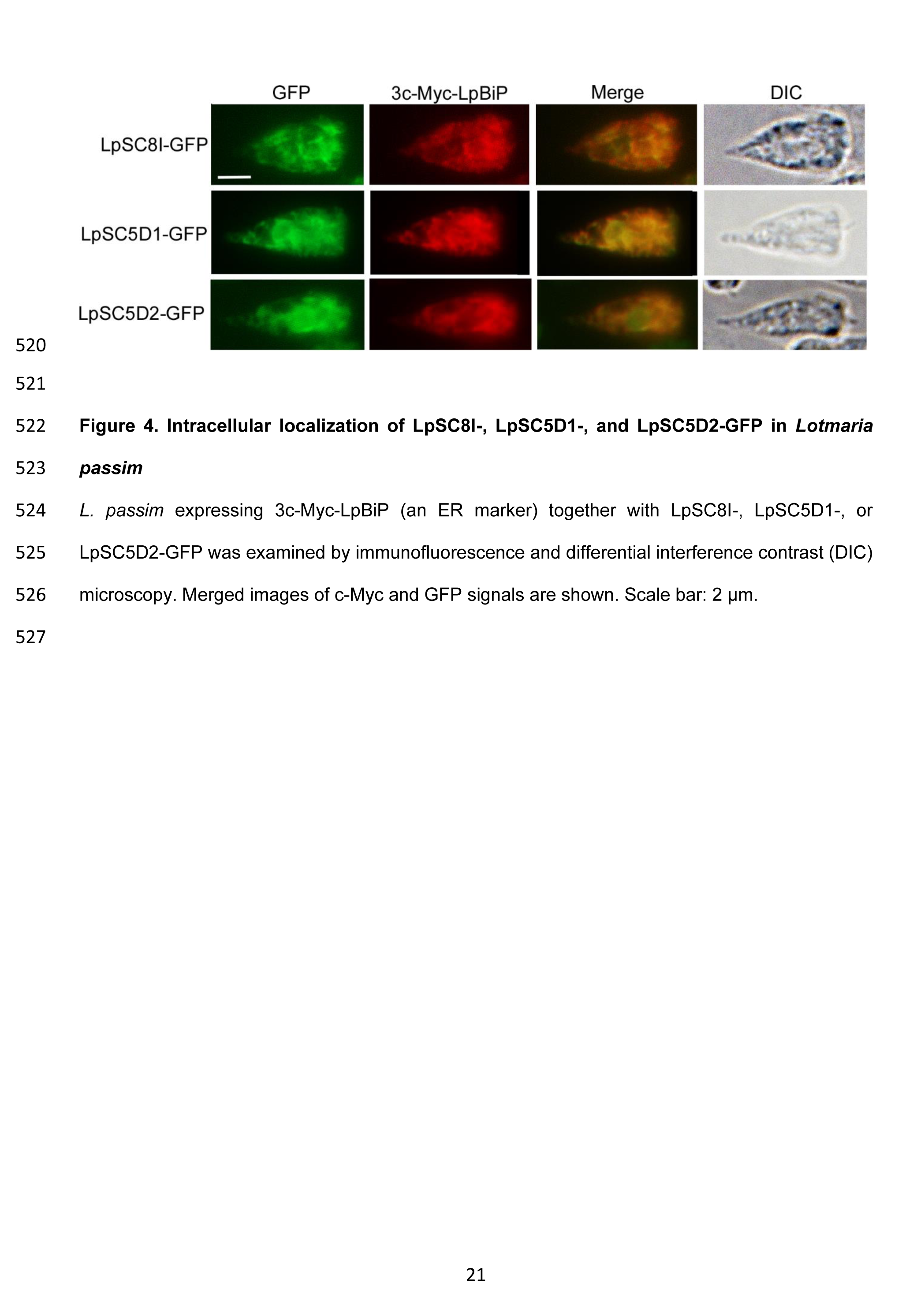
**Intracellular localization of LpSC8I-, LpSC5D1-, and LpSC5D2-GFP in *Lotmaria passim*** (A) *L. passim* expressing 3c-Myc-LpBiP (an ER marker) together with LpSC8I-, LpSC5D1-, or LpSC5D2-GFP was examined by immunofluorescence and differential interference contrast (DIC) microscopy. Merged images of c-Myc and GFP signals are shown. Scale bar: 2 μm.

**Figure 5.**
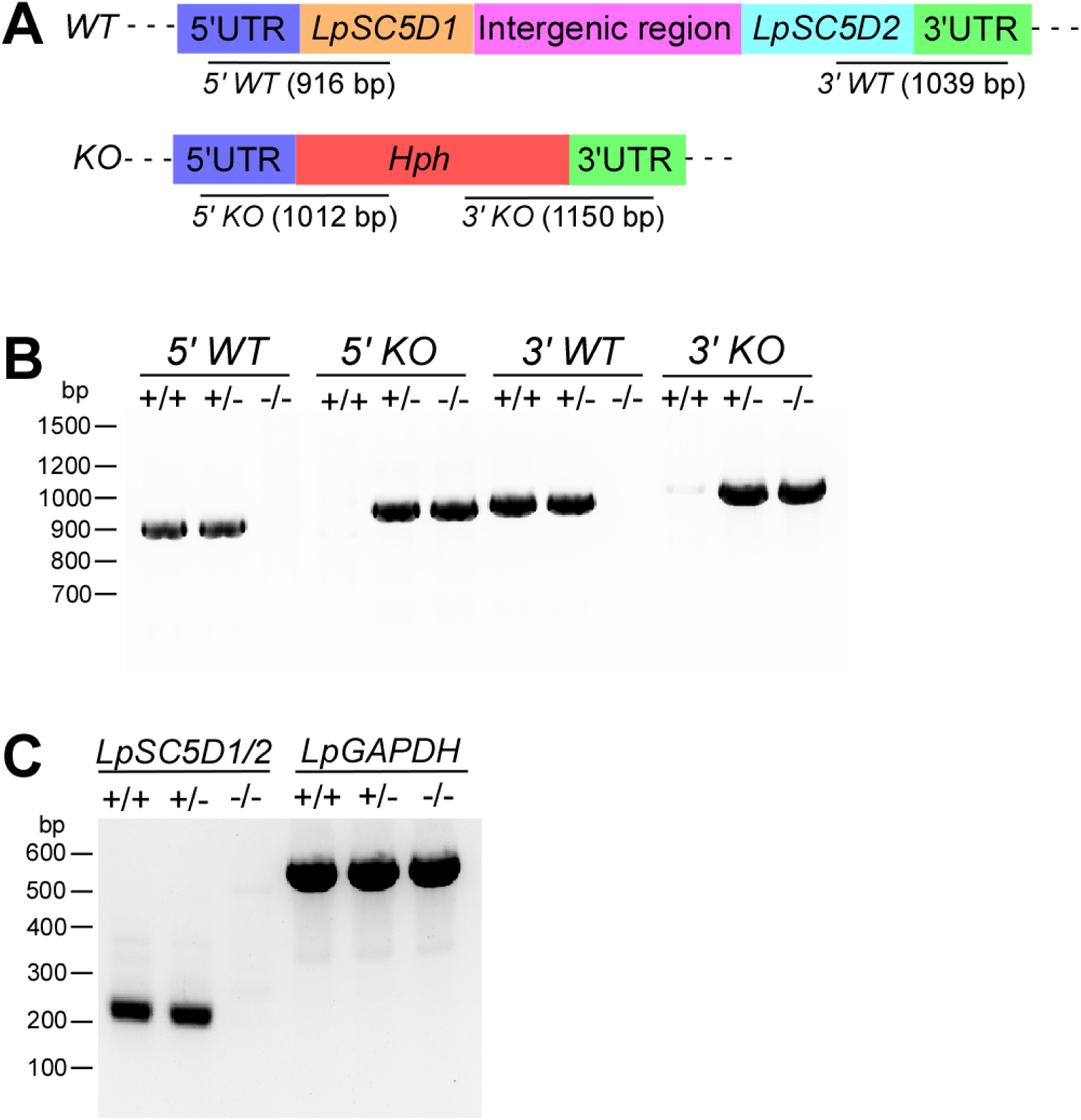
**CRISPR-mediated deletion of *LpSC5D*** (A) Schematic of WT and deleted (KO) *LpSC5D* alleles generated by CRISPR/Cas9-induced homology-directed repair. The 5′ and 3′ UTRs, ORFs, intergenic region, and *Hph* are color-coded. Expected PCR product sizes for detecting 5′ WT, 3′ WT, 5′ KO, and 3′ KO alleles (not to scale) are shown. (B) Genomic DNA from WT (+/+), heterozygous (+/–), and homozygous (–/–) *LpSC5D* mutants was analyzed by PCR for the 5′ WT, 5′ KO, 3′ WT, and 3′ KO alleles. Molecular weight markers are indicated on the left. (C) RT-PCR detection of *LpSC5D1/2* (*LpSC5D1* and *LpSC5D2*) and *LpGAPDH* mRNAs in WT, heterozygous, and homozygous *LpSC5D* mutants using a forward primer specific to the *L. passim* splice leader sequence. Molecular weight markers are shown on the left.

### Phenotypes of *LpSC5D*-deficient parasites

To investigate the role of LpSC5D, we complemented mutants with plasmid-based expression constructs carrying either *LpSC5D1* or *LpSC5D2*, together with a bleomycin resistance marker (*LpSC5D*+LpSC5D1 or *LpSC5D*+LpSC5D2). Growth assays showed that while mutants grew normally at 30 °C (Fig. 6A), they exhibited severely reduced growth at 20 °C (Fig. 6B). Thus, ergosterol is required for efficient growth at low temperature. During mid-log growth at 20 °C, wild-type parasites displayed elongated cell bodies with typical epimastigote morphology. In contrast, *LpSC5D*-deficient parasites had smaller, rounded cell bodies (Fig. 6C and D). Ectopic expression of either *LpSC5D1* or *LpSC5D2* rescued morphology but only partially growth (Fig. 6B–D). As in *Leishmania* spp. [22, 28], wild-type *L. passim* was sensitive to Amphotericin B (AmB), with growth strongly inhibited at 0.1 µg/mL at 30 °C. Intriguingly, *LpSC5D*-deficient parasites grew normally in the presence of AmB (Fig. 6E). Complemented strains (*LpSC5D*+LpSC5D1 and *LpSC5D*+LpSC5D2) grew more slowly in AmB than mutants, but eventually reached the same density after four days (Fig. 6E and F). AmB selection pressure likely causes parasites to lose episomal plasmid DNA encoding LpSC5D1 or LpSC5D2. Notably, during the 2-3 days, *LpSC5D*+LpSC5D1 grew significantly more slowly than *LpSC5D*+LpSC5D2 (Fig. 6F).

**Figure 6.**
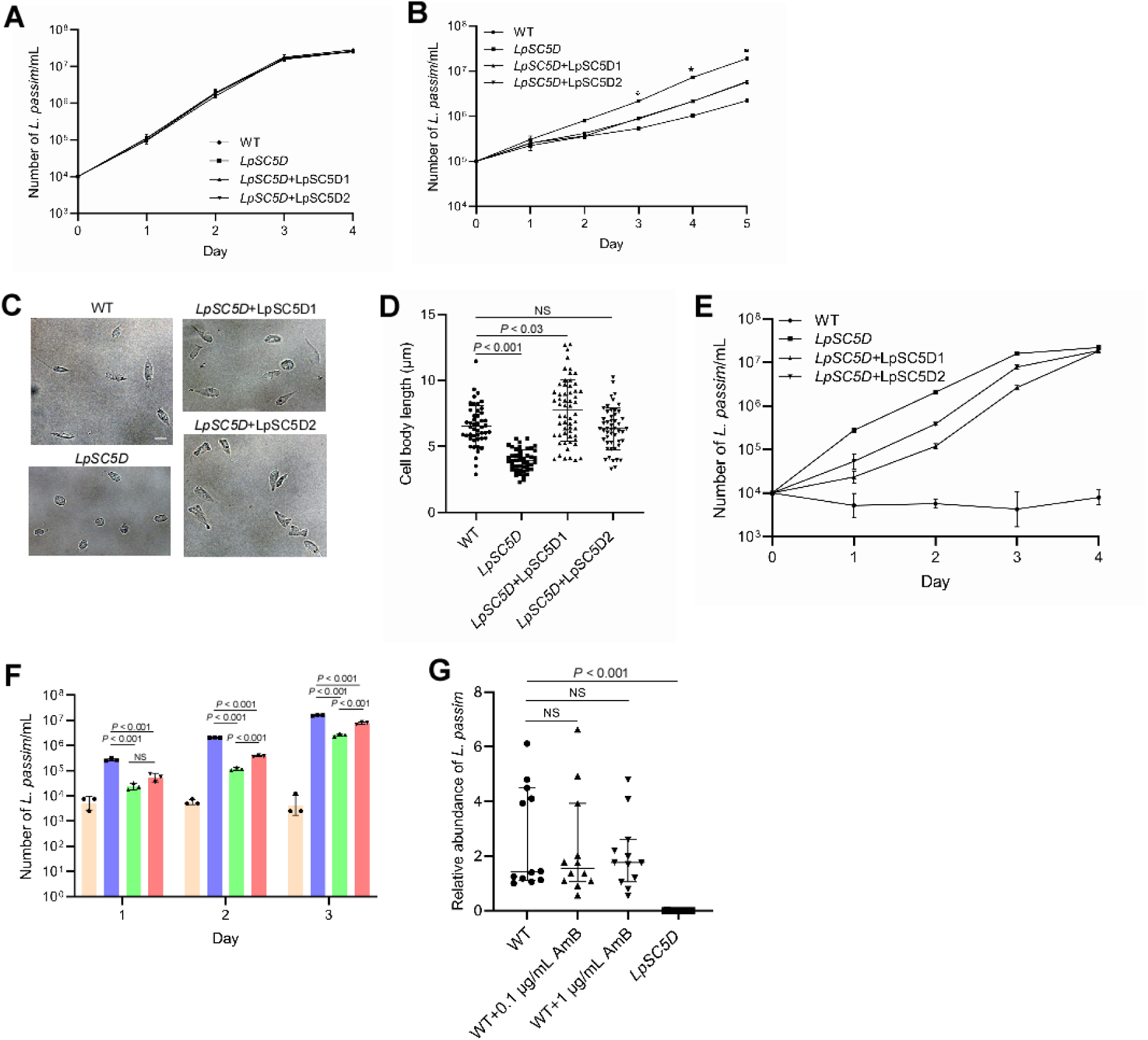
Phenotypes of *LpSC5D*-deficient mutants in culture and honey bees. (A) Growth of WT (circles), *LpSC5D* homozygous mutant (clone C2, squares), *LpSC5D*+*LpSC5D1* (triangles), and *LpSC5D*+*LpSC5D2* (inverted triangles) parasites at 30 °C over 4 days (biological replicates, n = 3). (B) Growth of the same strains at 20 °C over 5 days (biological replicates, n = 3). Statistically significant differences in growth between WT and *LpSC5D* mutants, WT and *LpSC5D*+*LpSC5D1/2*, and mutants vs. complemented strains are indicated (asterisks; day 3: *P* < 0.001, *P* < 0.001, *P* < 0.005; day 4: *P* < 0.001, *P* < 0.001, *P* < 0.001; day 5: *P* < 0.001, *P* < 0.001, *P*< 0.02; Tukey HSD test). (C) Morphology of WT, *LpSC5D* mutant, and complemented parasites. Scale bar: 5 µm. (D) Cell body length of WT (n = 47), *LpSC5D* mutant (n = 45), *LpSC5D*+*LpSC5D1* (n = 58), and *LpSC5D*+*LpSC5D2* (n = 45). Data are shown as medians with 95% CI. Significant differences were found between WT and *LpSC5D* mutants and between WT and *LpSC5D*+*LpSC5D1* (Steel test). NS: not significant. (E) Growth of WT, *LpSC5D* mutant, and complemented parasites with 0.1 µg/mL Amphotericin B (AmB) at 30 °C over 4 days (biological replicates, n = 3). (F) Cell density of WT (beige), *LpSC5D* mutant (blue), *LpSC5D*+*LpSC5D1* (green), and *LpSC5D*+*LpSC5D2* (red) with AmB across days 1–3. Statistical analysis by Tukey HSD test. (G) Relative abundance of *L. passim* in honey bees (n = 12) 14 days post-infection with WT, WT fed 50% sucrose containing 0.1 or 1 µg/mL AmB, or *LpSC5D* mutant parasites. Data were normalized to one WT-infected sample without AmB (set as 1) and are shown as medians with 95% CI. Statistical analysis by Steel test.

### LpSC5D is required for conversion of Ergosta-7,22-dienol to Ergosterol

GC-MS analysis revealed that wild-type *L. passim* produces ergosterol as its major sterol. In contrast, *LpSC5D*-deficient parasites predominantly accumulated ergosta-7,22-dienol (Fig. 7A), confirming that LpSC5D catalyzes its conversion to ergosterol (Fig. 3). Complemented parasites produced ergosterol as the major sterol, though significant amounts of ergosta-7,22-dienol remained. The ergosterol-to-ergosta-7,22-dienol ratio was higher in *LpSC5D*+LpSC5D1 than in *LpSC5D*+LpSC5D2 (Fig. 7B), consistent with the slower AmB growth of *LpSC5D*+LpSC5D1 (Fig. 6F). Assuming equal plasmid copy number, LpSC5D1 appears more active than LpSC5D2.

**Figure 7.**
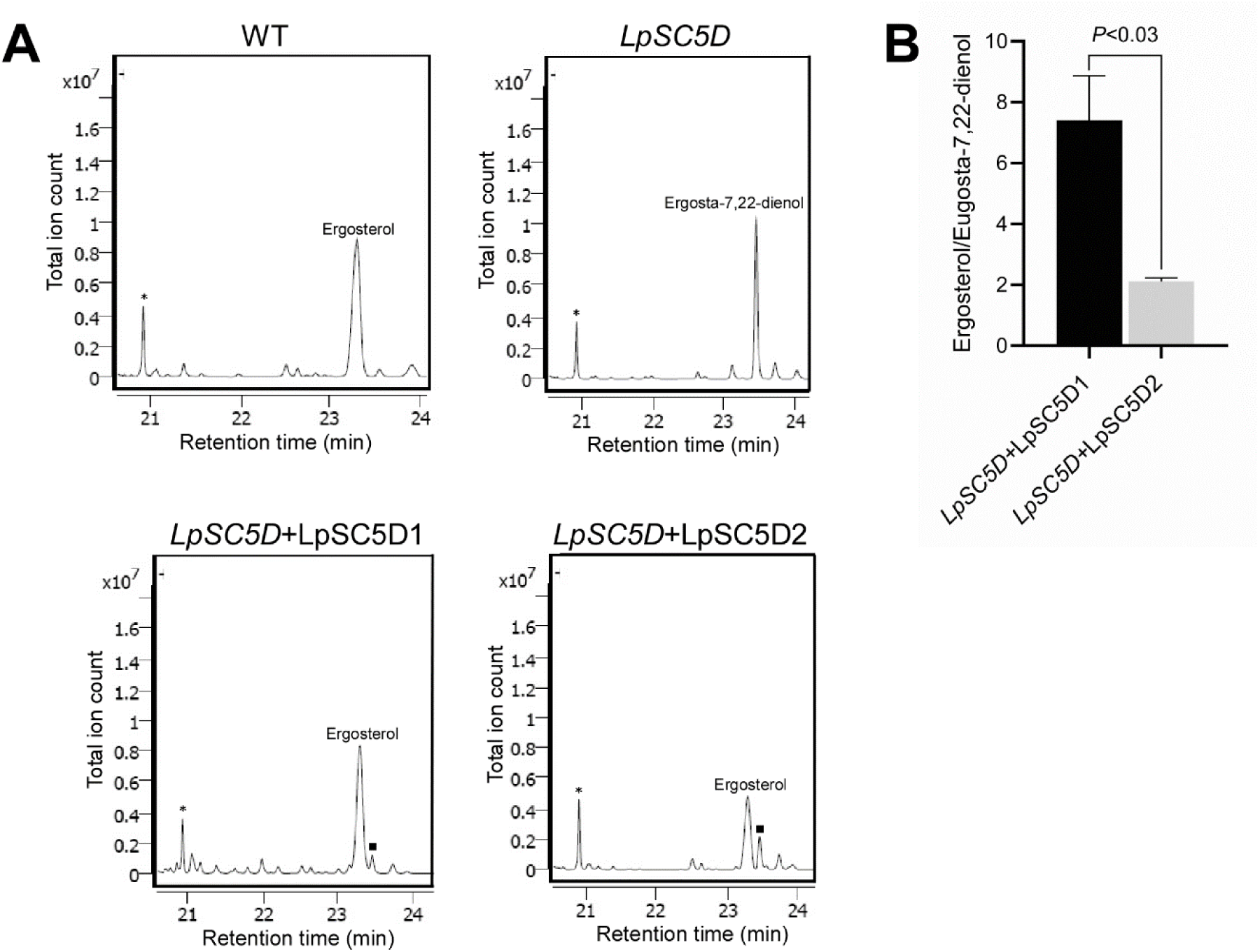
Accumulation of ergosta-7,22-dienol in *LpSC5D* mutants and restoration of ergosterol synthesis in rescued parasites. (A) Partial GC-MS spectra of total sterols from WT, *LpSC5D* mutant, *LpSC5D*+*LpSC5D1*, and *LpSC5D*+*LpSC5D2* parasites. Cholesta-3,5-diene (retention time = 20.9 min) was added as an internal standard (asterisks). Analyses were performed three times, and percentages of ergosterol (retention time = 23.3 min) and ergosta-7,22-dienol (retention time = 23.5 min) were quantified. Peaks corresponding to ergosta-7,22-dienol in *LpSC5D*+*LpSC5D1* and *LpSC5D*+*LpSC5D2* parasites are marked by squares. (B) Ratios of ergosterol to ergosta-7,22-dienol in *LpSC5D*+*LpSC5D1* and *LpSC5D*+*LpSC5D2* parasites. Statistical analysis was performed using Welch’s *t*-test.

### Ergosterol is essential for honey bee gut infection by *L. passim*

We next tested infectivity in honey bees. Fourteen days after oral infection, *LpSC5D*-deficient parasites showed dramatically reduced colonization of the gut compared to wild-type parasites (Fig. 6G). This demonstrates that ergosterol is required for efficient honey bee gut infection. Feeding bees with sucrose containing 0.1 or 1 µg/mL AmB did not inhibit wild-type infection, likely due to drug inactivation in the gut environment (Fig. 6G). *Leishmania mexicana* and *L. major* lacking sterol C14-α-demethylase (CYP51), sterol C24-methyltransferase (SMT1/2), or SC5D were previously generated [21–23, 29, 30]; however, none of these mutants have been tested for transmission competence in their sand fly vectors. These results demonstrate that ergosterol biosynthesis is critical for *L. passim* colonization of the honey bee gut. Our findings support a conserved role of sterol metabolism in kinetoplastid parasite biology, shaped in part by horizontally acquired genes.

## Discussion

Our analyses demonstrate that HGT has played a selective yet consequential role in shaping the metabolic repertoire of trypanosomatids. The stark contrast between the numerous HGT candidates identified in the free-living kinetoplastid *B. saltans* and the reduced sets in parasitic species supports the hypothesis that the transition to parasitism was accompanied by extensive gene loss. Only a subset of horizontally acquired genes was retained, likely under strong selective pressure. Dollo parsimony analysis suggests that most HGT-derived genes were discarded early in kinetoplastid evolution, but several families persisted and underwent lineage-specific expansions. These findings imply that episodic HGT events contributed not only to early metabolic innovation but also to diversification within the parasitic lineages.

An example of such innovation is the recruitment of enzymes involved in redox balance and nitrogen metabolism under anaerobic conditions. Aspartate ammonia ligase, which is absent in *B. saltans* but conserved in trypanosomatids, likely originated via HGT. Together with aspartate ammonia lyase, this enzyme may form a nitrogen–redox loop that mitigates ammonia accumulation while supporting succinate fermentation. This adaptation is particularly relevant in the oxygen-poor environments of insect guts, where maintaining redox balance is essential for parasite survival and transmission. Thus, HGT-derived enzymes provided immediate ecological advantages that facilitated colonization of new hosts.

Our results reveal that early steps of ergosterol biosynthesis in trypanosomatids are mosaic in origin, incorporating enzymes of bacterial and archaeal ancestry. Functional analyses in *L. passim* highlight the centrality of this pathway: LpSC8I is essential for parasite viability, while LpSC5D, although dispensable for *in vitro* proliferation, is required for ergosterol synthesis, low temperature tolerance, and drug susceptibility. Strikingly, loss of ergosterol biosynthesis abolished parasite colonization of the honey bee gut, underscoring that sterol composition is critical not only for membrane integrity but also for successful host infection. These results provide a direct link between horizontally acquired enzymes, a central metabolic pathway, and parasitic fitness.

This study illustrates how selective retention of HGT-derived genes has shaped the evolution of trypanosomatids. By integrating comparative genomics with functional assays, we show that horizontally acquired genes were incorporated into core metabolic network which is critical for growth at low temperature and host colonization. Together, these findings highlight the dual role of HGT in both metabolic innovation and the emergence of lineage-specific adaptations that underlie parasitic success.

### Materials and Methods Identification of HGT candidates

We employed the DarkHorse [24, 31] algorithm to predict potential HGTs in 12 trypanosomatid species and *B. saltans*. Protein sequences of *L. passim* were annotated using BRAKER [32, 33], while sequences of other species were obtained from TriTrypDB [34]. To run the DarkHorse analysis, the following steps were performed. First, a BLASTP search was conducted using DIAMOND v2.0.14 [35] against the DarkHorse HGT candidate database for all species, with an e-value cutoff of 1e-5. The top 100 most similar homologs were retained for further analysis. Next, an exclude_list file was generated to remove self-matches from the BLAST results, incorporating taxonomic information from the GenBank Taxonomy database. The darkhorse.pl script was then used to predict HGT events, identifying genes exhibiting a LPI below the defined threshold.

### Analyzing gain and loss of HGT-derived gene families

Orthologous groups of the identified HGT candidates were inferred across the 13 kinetoplastid species using OrthoFinder v2.5.4 [36, 37], allowing classification of conserved and lineage-specific gene families. To establish a reliable evolutionary framework, a species tree was constructed using core conserved genes (non-HGT genes), ensuring an unbiased representation of phylogenetic relationships. Gene family gains and losses were reconstructed along the species tree using COUNT [38], implementing Dollo parsimony (assuming single-origin gains). This approach enabled inference of gene family gain and loss patterns across kinetoplastida lineages. The final gene gains and losses were visualized by annotating the species tree using iTOL (Interactive Tree of Life) [39] for comprehensive evolutionary interpretation.

### Phylogenetic analysis of HGT candidates

For each target gene, the top 250 homologs were identified via NCBI BLASTP search. When top hits were bacterial or archaeal proteins, homologs from diverse eukaryotic lineages—*Homo sapiens, Branchiostoma floridae, Nematostella vectensis, Arabidopsis thaliana, Oryza sativa, Chlamydomonas reinhardtii, Dictyostellum discoideum, Acanthamoeba castellanii, Guillardia theta, Saccharomyces cerevisiae, Agaricus bisporus, Emiliania huxleyi, Tetrahymena thermophila, and Phytophthora infestans*—were retrieved. These species represent Opisthokonta, Amoebozoa, Excavata, SAR, and Archaeplastida [40]. If no homologs were found in these species, related species were used, such as *Chara braunii, Liolophura sinensis, Hydractinia symbiolongicarpus*, and *Patella vulgata*. Multiple sequence alignments were performed using MUSCLE v5 [41], followed by trimming poorly aligned regions with trimAl [42] to improve alignment quality. Processed alignments were used to construct maximum likelihood (ML) trees using IQ-TREE2 [43], with optimal substitution models automatically selected. One thousand bootstrap replicates were performed to assess nodal support, and resulting trees were visualized using iTOL.

### Cellular localization of 3c-Myc-LpBiP, LpSC8I-GFP, LpSC5D1-GFP, and LpSC5D2-GFP in *L. passim*

To express triple c-Myc-tagged LpBiP (3c-Myc-LpBiP), the full open reading frame (ORF) of *LpBiP* was amplified by PCR using KOD-FX DNA polymerase (TOYOBO), *L. passim* genomic DNA, and primers 5-XhoI-LpBiP/3-XhoI-stop-LpBiP. The PCR product was digested with XhoI and cloned into a vector carrying triple c-Myc epitopes [44], digested with the same enzyme. To construct LpSC8I-, LpSC5D1-, and LpSC5D2-GFP vectors, the ORFs were amplified using primer pairs LpSC8I-5-XbaI/LpSC8I-3-XbaI, LpSC5D1-5-XbaI/LpSC5D-3-XbaI, and LpSC5D2-5-XbaI/LpSC5D-3-XbaI. PCR products were digested with XbaI and cloned into the XbaI site of pTrex-n-eGFP plasmid [45] (Addgene #62544). All sequences are listed in Additional file 1. Actively growing *L. passim* cells (4 × 10⁷) were washed twice with PBS and resuspended in 0.4 mL Cytomix buffer (without EDTA) containing 20 mM KCl, 0.15 mM CaCl₂, 10 mM K₂HPO₄, 25 mM HEPES, and 5 mM MgCl ₂ (pH 7.6). Electroporation was performed twice at 1-minute intervals with 10 μg plasmid DNA using a Gene Pulser X cell electroporator (Bio-Rad) with a 2-mm gap cuvette, voltage 1.5 kV, capacitance 25 μF, and infinite resistance. Transfected parasites were cultured in 4 mL modified FPFB medium [46], and blasticidin (50 μg/mL) and G418 (200 μg/mL) were added after 24 hours to select drug-resistant clones.

Immunofluorescence detection was performed by washing and mounting parasites on poly-L-lysine-coated 8-well chamber slides, fixing with 4% paraformaldehyde, permeabilizing with 0.1% Triton X-100 in PBS (PT), and blocking with PT containing 5% normal goat serum (PTG). Samples were incubated overnight at 4 °C with rabbit anti-GFP polyclonal antibody (1:1000, Proteintech) and CoraLite594-conjugated Myc tag monoclonal antibody (1:500, Proteintech) in PTG. After five washes with PT, samples were incubated with FITC-conjugated anti-rabbit IgG (Beyotime) for 2 hours at room temperature, washed, stained with DAPI (1 µg/mL) for 5 minutes, and observed under a Nikon Eclipse Ni-U fluorescence microscope with 200 ms exposure.

### Disruption of *LpSC8I* and *LpSC5D* genes by CRISPR

Complementary oligonucleotides for sgRNA sequences (LpSC8IgRNA378F/R and LpSC5DgRNA688RF/RR) were phosphorylated by T4 polynucleotide kinase (TAKARA), annealed, and cloned into BbsI-digested pSPneogRNAH vector [47] (ADDGENE: # 63556). The LpSC5D gRNA targets both *LpSC5D1* and *LpSC5D2*. *L. passim* expressing Cas9 [27] was electroporated with 10 μg plasmid DNA, and transformants were selected with blasticidin and G418. Donor DNA for *LpSC8I* consisted of 5’UTR (LpSC8I-5’UTR-F and LpSC8I-5’UTR-R), *Hph* ORF (LpSC8I-Hph-F and LpSC8I-Hph-R), and 3’ UTR (LpSC8I-3’UTR-F and LpSC8I-3’UTR-R) fragments. They were fused and cloned into the EcoRV site of pBluescript II SK(+). For *LpSC5D* genes, 5’UTR of *LpSC5D1*, *Hph* ORF, and 3’UTR of *LpSC5D2* were fused to replace ORFs of both *LpSC5D1* and *LpSC5D2* by *Hph*. The linearized plasmid DNA (20 μg) digested with EcoRI (for *LpSC8I* donor DNA) or BamHI (for *LpSC5D* donor DNA) was used for electroporation of *L. passim* expressing both Cas9 and *LpSC8I* or *LpSC5D* gRNA, as described above. *L. passim* resistant to blasticidin, G418, and hygromycin was selected and cloned by serial dilutions in a 96-well plate. The genotype of each clone was first determined by the detection of 5’ wild-type (WT) and deleted (KO) alleles for *LpSC8I* or *LpSC5D1* by PCR. After identifying heterozygous (+/-) and homozygous (-/-) KO clones of *LpSC5D*, their 5’WT (LpSC5D1-5’UTR-Up and LpSC5D1-63R), 5’KO (LpSC5D1-5’UTR-Up and Hyg-159R), 3’WT (LpSC5D2-1782F and LpSC5D2-3’UTR-Down), and 3’KO (Hyg-846F and LpSC5D2-3’UTR-Down) alleles were confirmed by PCR. For *LpSC8I*, only heterozygous KO clones were identified by testing the 5’WT (LpSC8I-5’UTR-Up and LpSC8I-32R) and KO alleles (LpSC8I-5’UTR-Up and Hyg-159R).

### RT-PCR

Total RNA from WT, *LpSC5D* heterozygous, and homozygous mutants was extracted using TRIzol (Sigma-Aldrich). Reverse transcription of 0.2 μg RNA was performed with ReverTra Ace (TOYOBO) and random primers, followed by PCR with GoTaq Green Master Mix (Promega). Both *LpSC5D1* and *LpSC5D2* mRNAs were detected using nested PCR, with primers LpSL-F-1st/LpSC5D-250R for first round and LpSL-F-2nd/LpSC5D1-63R for second round. Two reverse primers are complementary to both *LpSC5D1* and *LpSC5D2* mRNAs. *LpGAPDH* mRNA was detected with LpSL-F and LpGAPDH-R primers.

### Cultute and cell body length measurement of *L. passim*

Plasmid DNA expressing LpSC5D1 or LpSC5D2 was constructed by amplifying ORFs, using LpSC5D1-5-XbaI and LpSC5D-stop-HindIII or LpSC5D2-5-XbaI and LpSC5D-stop-HindIII primers. The plasmid DNA to express the corresponding GFP fusion protein was used as a template. The resulting PCR product was digested with XbaI and HindIII followed by subcloning into the same restriction enzyme sites of the tdTomato/pTREX-b plasmid DNA [48] (ADDGENE: #68709) with bleomycin resistance gene. *LpSC5D*-deficient parasites were electroporated with above plasmid DNA and the parasites expressing LpSC5D1 (*LpSC5D*+LpSC5D1) or LpSC5D2 (*LpSC5D*+LpSC5D2) were selected with hygromycin and zeocin.

WT, *LpSC5D*-deficient, *LpSC5D*+LpSC5D1, and *LpSC5D*+LpSC5D2 parasites were inoculated into the culture medium at 10⁴ cells/mL at 30 °C. Cell counts were performed daily for 4 days using a hemocytometer. AmB was added at 0.1 µg/mL when necessary. For growth at 20 °C, parasites were inoculated at 10⁵ cells/mL and counted for 5 days. Cell body length was measured at day 4 using Image-J. Two independent *LpSC5D*-deficient clones (A1 and C2) were phenotypically identical; C2 was used for further experiments.

### GC-MS analysis of sterol in *L. passim*

Parasites were cultured to 10⁷ cells/mL in triplicate with 10 mL culture medium. Zeocin (50 µg/mL) was added to maintain plasmids in *LpSC5D*+LpSC5D1 and *LpSC5D*+LpSC5D2 parasites. Cells were collected, washed with 5 mL PBS, and extracted with 3 mL chloroform:methanol (2:1), with 20 µg cholesta-3,5-diene as internal control. 0.6 mL of 0.9% NaCl was added to separate phases; the lower phase was dried and suspended in 0.3 mL methanol. Electron impact GC/MS analysis of sterol was carried out using a Thermo Scientific ISQ single-stage quadrupole mass spectrometer coupled with a Trace GC system, operated through Thermo Xcalibur 2.1 software. Extracts (1 mL) were injected in splitless mode and separated on a Phenomenex ZB-50 column (15 m × 0.32 mm i.d., 0.5 µm film thickness). The GC oven was programmed as follows: an initial temperature of 100 °C held for 2 min, ramped to 200 °C at 50 °C/min, then increased to 300 °C at 10 °C/min, with a final hold at 300 °C for 10 min. The injector and transfer line temperatures were maintained at 280 °C, and the ion source at 220 °C. Mass spectra were acquired in full-scan mode (m/z 50–500) or as total ion current chromatograms, at a rate of one scan every 0.2 sec. Electron ionization was performed at 70 eV.

### Honey bee infection

Log-phase parasites (5 × 10⁵ cells/mL) were washed and suspended in 10% sucrose/PBS at 5 × 10⁴ cells/μL. Newly emerged honey bees were starved for 2–3 hours, then 20 bees were fed 2 μL solution containing either WT or *LpSC5D*-deficient parasites (10⁵ cells in total). WT-infected bees were fed 50% sucrose or sucrose containing 0.1 or 1 μg/mL AmB. Bees were maintained at 33 °C for 14 days and frozen at −80 °C. Four bees from each of three repeats (12 total per group) were analyzed. Genomic DNA from the whole abdomen of individual bee was extracted using DNAzol (Thermo-Fisher). *L. passim* abundance was quantified by qPCR with LpITS2-F/R primers targeting the internal transcribed spacer region 2 in the ribosomal RNA gene; honey bee AmHsTRPA was the reference gene with AmHsTRPA-F/R primers [49]. Relative abundance was calculated using the ΔCt method, setting one WT sample fed 50% sucrose as 1. Statistical analysis was conducted with the Brunner-Munzel test. All primer sequences are listed in Additional file 2.

## Acknowledgments

We thank Jinji Lake Double Hundred Talents Programme for financial support.

