## Additional file 1 for "Essential roles of HGT-derived sterol biosynthesis in trypanosomatid growth and parasitism"

>LpBiP

ATGGCAGTGAGGGATCGTCTTGTGCTGCTGGCGGTGTGCCTCGTGTCTGCACTGCTCATCGCAGCTGCGG

TCGCTGCACCGGACGGCAGTGGCAAGGTGGAGCCCCCGTGCATCGGCGTCGACCTCGGTACGACCTACTC

TGTCGCTGCCGTTTGGCAGAAGGGTGAGGTGCACATCATCACCAATGAAATGGGCAACCGCATCACCCCG

TCCGTTGTCGCCTTTACAGAGACGGAGCGGTTGGTCGGCGATGGTGCGAAGAACCAGCTCCCGCAGAACC

CGGAGAACACCATCTACGCCATCAAACGCCTGATTGGTCGTAAGTTCGCTGACCCAACGGTGCAGAACGA

CAAGAAGCTGCTGTCGTACAAGATTATCAGCGACAAGGCGGGCAAACCTCTCGTGCAGGTCACTGTGAAC

GGCGCCAAGAAGGAGTTCACGCCGGAGGAGGTGAGCGCGATGGTGCTGCAGAAAATGAAGGACATCTCGG

AGACCTTCCTTGGCGAGAAGGTGAAGAACGCCGTGGTGACGGTGCCCGCCTACTTCAACGACGCCCAGCG

CCAGGCGACGAAGGACGCCGGCAAGATTGCGGGCCTCAACGTGGTGCGCATCATCAATGAGCCGACGGCG

GCCGCCATCGCCTACGGCTTGAACAAGGCGGGGGAGAAGAACATCCTTGTCTTTGATCTGGGTGGCGGCA

CGTTCGATGTGTCGCTGCTGACCATCGACGAGGGCTTCTTTGAGGTGGTGGCGACGAACGGTGACACCCA

CCTCGGTGGCGAGGACTTCGATAACAGCATGATGAAGTTCTTCGTCGACGGCCTCAAGCGCAAGCAGAAC

ATCGATATTTCAAACGACCAGAAGGCGCTGGCGCGTCTGCGCAAGGCGTGCGAGGCGGCGAAGCGTCAGC

TCTCGTCGCACCCCGAGGCGCGTGTGGAGGTGGACAGCCTCGTAGAGGGCCACGACTTCAGCGAGAAGAT

TACGCGTGCCAAGTTTGAGGAGCTGAATATGGGCATGTTCAAGAACACCCTAATCCCGGTGCAGAAGGTG

CTGGAGGATGCGAAGCTGAAGAAGAGCGACATCGACGAGATCGTCCTCGTGGGTGGTTCGACCCGCATTC

CGAAAGTGCAGCAGCTGATCAAGGACTTCTTCGGCGGCAAGGAGCCGAACAAGGGCATCAACCCGGACGA

GGCCGTGGCGTATGGTGCGGCGGTGCAGGCGGCTGTGCTGATGGGTGAAAGCGAGGTCGGCGGCAAGGTC

GTTCTTGTTGATGTGATTCCCCTTTCCCTCGGCATCGAGACCGTCGGCGGCGTGATGACGAAGCTGATCG

AGCGCAACACGCAGATCCCGACCAAGAAGAGCCAGGTCTTCTCCACCTACCAGGACAACCAGCCCGGCGT

GCTCATCCAGGTTTTCGAAGGCGAACGGCAGATGACAAAGGACAATCGCCTACTAGGCAAGTTCGAGCTC

TCCGGCATCCCGCCGGCGCCGCGCGGAGTTCCACAGATCGAGGTCGCCTTCGACGTGGACGAGAACAGCA

TTCTGCAGGTGTCGGCGAGTGACAAGTCGTCTGGCAAGCGGGAGGAGATCACCATCACGAACGACAAGGG

TCGCCTGAGCGATGCGGAGATCCAGGCAATGGTGGAGGAAGCCGCGCAATTCGCTGAGGAGGACCGCAAG

GTGCGGGAGCGCGTGGAGGCGAAGAACTCGCTGGAGAGCATCGCGTACTCCCTGCGCAACCAGATCAACG

ACAAGGAGAAGCTTGGTGACAAGCTGGACGCGGACGATAAGAAGGCGATTGAGGCTGCCGTGCAGGTGGC

GCTCGATTTCGTCGACGAGAACCCGAACGCGGACCGCGAGGAGTTCGAGGAGGCGCGCGAGCAGCTGCAG

AAGGTAACGAATCCGATTATTCAGAAGGTGTACCAGGCTGCTGGCGGTGCCGCTGGTGAGGAGCCGGACG

CGATGGACGACTTGTAA

>LpSC8I

ATGCTCGGTTCTCGCTTCTCTGTGATGCTGGTAGCCCTCGTGGCGATCCTCATCGCGTTCTTCGTCTACGTGGACCAGCCCTCCAACTGGGTGTACGATCCGGCGCGCCTGCAGCAAATTGCGCAGCAGAGTATCGCCAACGCGAAGGCCGCCCATGGTGAGCACGCCACGGCAAAGCAGATCACCGACGAGACAATTCGGTTGATGCTCGAGGCGTACCCGCAGACGACGCGGAGCACCGGTCACTGGCTGTGGAACAACGCGGGCGGGGCGATGGGCTCCATGACGGTGCTGCACTGCTCCTTCTCGGAGTACATCATCATCTTCGGTACACCGGTCGGGACGGAGGGCCACACCGGGCGCTACTTCTGGGCCGAGGACTTCTTCAACATACTGGTGGGCGAGCAGTGGGCGGCGCTGCCTGGTGTGGCAGAGAGGGAGGTGTACCGCCCAGGTGATCAGCACGTCCTGCCGCGTGGCGTGGCGAAGCAGTACCGCATGCCGGACGAGTGCTGGGCTCTCGAGTATGCCCGTGGCAACATCGTCTCCATGCTCTTCTTTGGCTTCGCTGATATGCTGTCCTCGACGTTGGACGTGGTCACGACGTGGCATACGGTAGTGGAGAGTCTTGGCAACATGATCCCAAACCTGCTCGCTGGCAAGATTTAA

>LpSC5D1

ATGGACTTTGCCTTCAGGCTTTACGCCTCCGTCCTGCCGGTGGACAAGGACAAACTGACGCATCAGATGTTCATTTTTTGGCTGATCCTGACAACGGGTGGCACCTTCATGTACCTCTCCTTCGCTTCGCTGTCCTACAACATCTACTTCCGCCGCCTGAAGCAGCAGTTCTTCCCCAAGACGATCGACCCGGACAATACAACAGAACTACGACGGCAGGCCCTGCACGAGATCTGGATAGCGACGTGCTCCATCCCGTTCATGGCAGTACTGATGATGCCGGCTGCTGTCTTCTCGCACCGCGGCTACAGCAAAGTGTACTACAACATCTCCGATTACGGCTGGGCCTACTTCTTCCTCTCGCCCGTGCTGTTCTTCGCATTCACAGACTTCATGGTGTACTGCTTCCACCGTGGCTTGCATCACCCGATCATCTATAAGCACGTCCATAAGCTGCATCACACGTACAAGTTCACGACGCCCTTCTCCTCGCACGCCTTCAACCCGGTCGACGGGTTCGGTCAGGGTGTGCCGTACTACATCTTTGTTTATCTGTTCCCCCTGCACAATGTACTCTTCATGTGTCTTTTTGTGATGGTGAACTTCTGGACTATCTCGATTCACGACCAGGTGGACTTTGGCGGCCACTTTCTGAACACGACGGGCCACCACACGATCCACCACGAGCTGTTCAACTACGACTACGGGCAGTACACGACAGTGTGGGACCGCCTGGGTGGGACGTATCGTCCCGCGGAGCAGACGCACCAGATGACGACGCTGCTGCACGCGTGTGACCAGAATTACGTGGACCCTGTGTACGCGACGTACCACGACGAGAAGGGCTTCCTGGCTGGCCGCTTCAAGGAGGAGGCTGCAGCGCGTCACGTCAAGAAGGCTGCGTAG

>LpSC5D2

ATGGACTTTGCCTTTGACCTTTACACGTCTGTCCTGCCGGTGGACAAGGACAAACTGACGCATCAGATGTTCATTTTTTGGCTGATCCTGACAACGGGTGGCACCTTCATGTACCTCTCCTTCGCTTCGCTGTCCTACAACATCTACTTCCGCCGCCTGAAGCAGCAGTTCTTCCCCAAGACGATCGACCCGGACAATATGGCGGAGTTGTGGCGACAGGTCAAGCACGAGATCTGGATAGCGACGTGCTCCATCCCGTTCATGGCAGTACTGATGATGCCGGCTGCTGTCTTCTCGCACCGCGGCTACAGCAAAGTGTACTACAACATCTCCGATTACGGCTGGGCCTACTTCTTCCTCTCGCCCGTGCTGTTCTTCGCATTCACAGACTTCATGGTGTACTGCTTCCACCGTGGCTTGCATCACCCGATCATCTATAAGCACGTCCATAAGCTGCATCACACGTACAAGTTCACGACGCCCTTCTCCTCGCACGCCTTCAACCCGGTCGACGGCTTCGGTCAGGGTGTGCCGTACTTCATTTTCGTCTATCTTTTTCCTCTCCATCACCTTCTTTTCATCGCGCTCTTCATGTTGGTGAACTTCTGGACTATCTCGATTCACGACCAGGTGGACTTTGGCGGCCACTTTCTGAACACGACGGGCCACCACACGATCCACCACGAGCTGTTCAACTACGACTACGGGCAGTACACGACAGTGTGGGACCGCCTGGGTGGGACGTATCGTCCCGCGGAGCAGACGCACCAGATGACGACGCTGCTGCACGCGTGTGACCAGAATTACGTGGACCCTGTGTACGCGACGTACCACGACGAGAAGGGCTTCCTGGCTGGCCGCTTCAAGGAGGAGGCTGCAGCGCGTCACGTCAAGAAGGCTGCGTAG
